# Size-Dependent Expression of the Mitotic Activator Cdc25 as a Mechanism of Size Control in Fission Yeast

**DOI:** 10.1101/078592

**Authors:** Daniel Keifenheim, Xi-Ming Sun, Edridge D'Souza, Makoto J. Ohira, Mira Magner, Michael B. Mayhew, Marguerat Samuel, Nicholas Rhind

**Affiliations:** Department of Biochemistry and Molecular Pharmacology, University of Massachusetts Medical School, Worcester MA 01605 USA; MRC London Institute of Medical Sciences (LMS), London W12 0NN UK; Institute of Clinical Sciences (ICS), Faculty of Medicine, Imperial College London, London W12 0NN UK; Lawrence Livermore National Laboratory, Livermore CA 94550 USA

**Keywords:** Cell Size, Cdc25, *S. pombe*, Unstable Activator, Excess Mitotic Delay

## Abstract

Proper cell size is essential for cellular function (Hall et al., 2004). Nonetheless, despite more than 100 years of work on the subject, the mechanisms that maintain cell size homeostasis are largely mysterious (Marshall et al., 2012). Cells in growing populations maintain cell size within a narrow range by coordinating growth and division. Bacterial and eukaryotic cells both demonstrate homeostatic size control, which maintains population-level variation in cell size within a certain range, and returns the population average to that range if it is perturbed (Marshall et al., 2012; Turner et al., 2012; Amodeo and Skotheim, 2015). Recent work has proposed two different strategies for size control: budding yeast has been proposed to use an inhibitor-dilution strategy to regulate size at the G1/S transition (Schmoller et al., 2015), while bacteria appear to use an adder strategy, in which a fixed amount of growth each generation causes cell size to converge on a stable average, a mechanism also suggested for budding yeast (Campos et al., 2014; Jun and Taheri-Araghi, 2015; Taheri-Araghi et al., 2015; Tanouchi et al., 2015; Soifer et al., 2016). Here we present evidence that cell size in the fission yeast *Schizosaccharomyces pombe* is regulated by a third strategy: the size dependent expression of the mitotic activator Cdc25. The *cdc25* transcript levels are regulated such that smaller cells express less Cdc25 and larger cells express more Cdc25, creating an increasing concentration of Cdc25 as cell grow and providing a mechanism for cell to trigger cell division when they reach a threshold concentration of Cdc25. Since regulation of mitotic entry by Cdc25 is well conserved, this mechanism may provide a wide spread solution to the problem of size control in eukaryotes.

## Results and Discussion

Size control in fission yeast is achieved by coordinating the timing of mitotic entry with cell size (Fantes, 1977). A number of hypotheses have been proposed for how fission yeast could measure their cell size. One elegant idea was that a gradient of the Pom1 mitotic inhibitor, which is concentrated at the cell poles and diffuses towards the cell center, might indirectly inhibit mitosis by phosphorylating key mitotic regulator at the cell midzone until the cell grew long enough that the concentration of the Pom1 gradient at the midzone dropped below a critical threshold, allowing mitotic entry (Martin and Berthelot-Grosjean, 2009; Moseley et al., 2009; Hachet et al., 2011). However, subsequent analysis showed despite the size-dependent gradient of Pom1, its level at the midzone does not correlate with mitotic regulation and, furthermore, its function is not required for cell-size regulation (Wood and Nurse, 2013; Bhatia et al., 2014). A more recent proposal suggests that Cdr2, the target of Pom1 inhibition, increases in concentration at the midzone as cells grow, and that mitosis is triggered when the concentration of Cdr2, a positive regulator of mitosis, reaches a certain threshold (Pan et al., 2014). However, the fact that cell lacking Cdr2 are viable and maintain size homeostasis, albeit at a larger size (Young and Fantes, 1987), indicates that there must be other mechanisms of cell size control in fission yeast.

The cell cycle machinery that regulates the entry into mitosis in fission yeast and other eukaryotes is well understood (Morgan, 2006), so we looked there for other potential regulators of cell size. Cdc25—the tyrosine phosphatase that dephosphorylates tyrosine 15 (Y15) of Cdc2, the catalytic subunit of the fission yeast CDK—is an attractive candidate. Y15 dephosphorylation of Cdc2 is the rate-limiting step for entry into mitosis (Gould and Nurse, 1989) and Cdc25 has been proposed to be involved in cell-size regulation (Moreno et al., 1990; Novak and Tyson, 1993). Its activity is balanced by Wee1, which phosphorylates Cdc2-Y15 (Russell and Nurse, 1987). We hypothesized that Cdc25, expressed in a size-dependent manner, would trigger entry into mitosis only when it reaches a certain threshold, ensuring that small cells stay in G2 and only sufficiently large cells enter mitosis. Specifically, we propose that the concentration of Cdc25 increases linearly with size. The amount of Cdc25 in the cell, which is the concentration times the cell size, would thus increase as the square of size. Such protein expression is unusual, since most proteins maintain a constant concentration as cells grow (Newman et al., 2006; Zhurinsky et al., 2010; Marguerat and Bahler, 2012; Marguerat et al., 2012).

To test our hypothesis, we measured the relative concentrations of Cdc25 in synchronous cultures, using the metabolic protein Ade4, which maintains a constant concentration independent of cell size (Figure S1), as an internal control. During G2, the concentration of Cdc25 increases about 2 fold (Figure 1A), consistent with our hypothesis and previous results (Moreno et al., 1990). In contrast Wee1, assayed in the same manner, maintains a relatively constant concentration during G2, as previously observed (Aligue et al., 1997), leading to an increasing Cdc25/Wee1 ratio as cells increase in size (Figures 1A and 1B). Both Cdc25 and Wee1 are unstable in G1 (Creanor and Mitchison, 1996; Aligue et al., 1997; Wolfe and Gould, 2004), resetting the system for the next G2.

**Figure 1:**
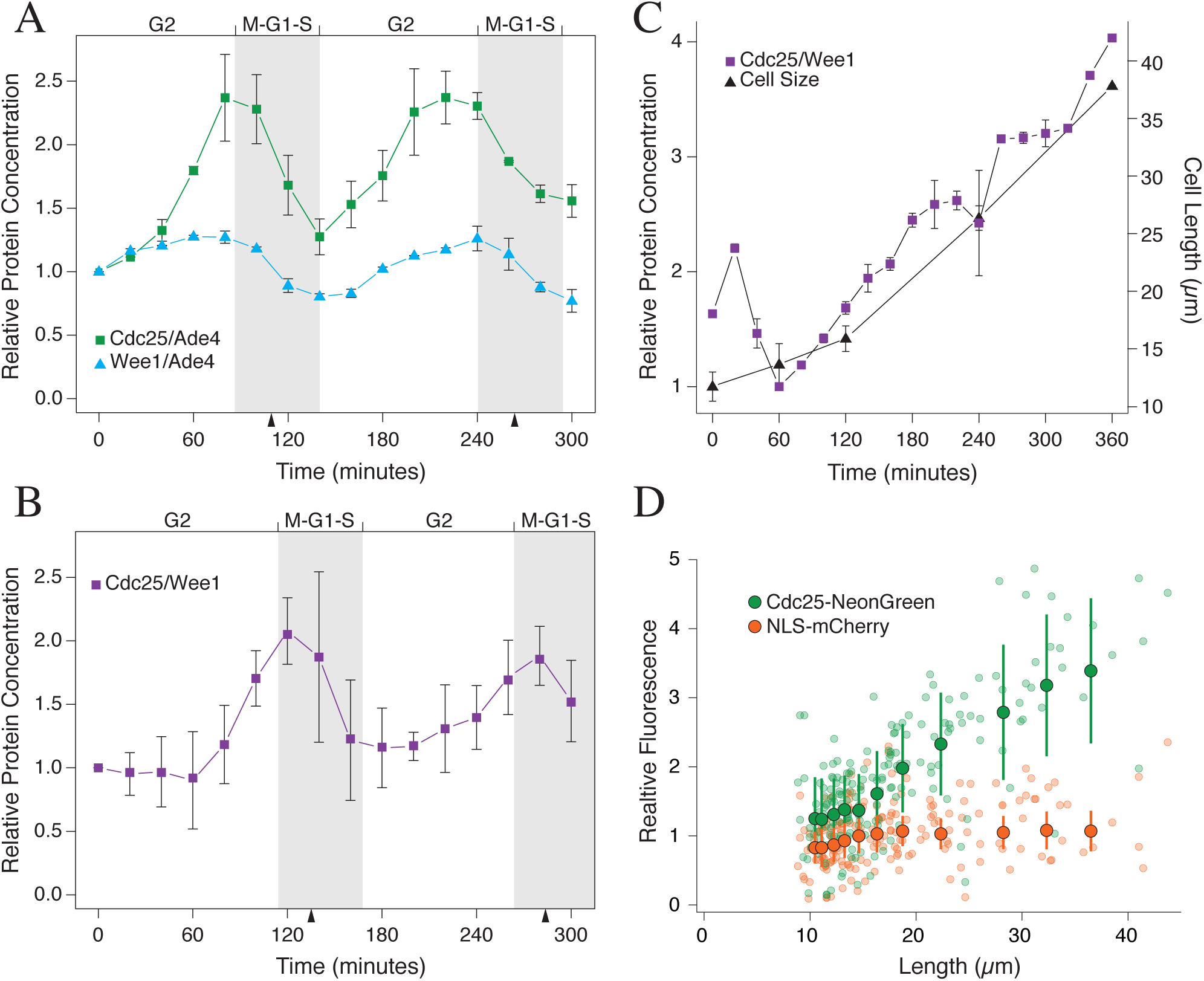
Cdc25 protein is expressed in proportion to cell size. **(A) Cdc25 protein concentration doubles during G2.** Cells expressing Cdc25-Rluc and Ade4-Bluc from their endogenous loci (yFS874) were elutriation synchronized in early G2 and followed through two synchronous cell cycles. Samples were taken every 20 minutes for luciferase quantitation and examined microscopically for septation. As a comparison, cells expressing Wee1-Rluc and Ade4-Bluc (yFS810) were similarly assayed. The midpoint of septation for each cycle is marked with an arrowhead and the inferred M-G1-S phases of the cycles are indicated in gray. The mean and standard error of the Ade4-nomalized Cdc25 and Wee1 signal, relative to time 0. n = 3 **(B) The Cdc25/Wee1 protein ratio doubles during G2.** Cells expressing Cdc25-Bluc and Wee1-Rluc (yFS870) were assayed as in **a**. The mean and standard error of the Cdc25/Wee1 signal, relative to time 0. n = 3 **(C) The Cdc25/Wee1 protein ratio increases linearly with cell size.** *cdc2-ts* cells expressing Cdc25-Bluc and Wee1-Rluc (yFS893) were shifted to the restrictive temperature of 35°C and sampled every 20 minutes. A transient increase in the Cdc25/Wee1 ratio was reproducibly seen after temperature shift. The mean and standard error of the Ade4-nomalized Cdc25 and Wee1 signal, relative to time 0. n = 3 **(D) Cdc25 protein concentration increases with cell size in individual cells.** *cdc2-ts* cells expressing Cdc25-NeonGreen and GST-NLS-mCherry (Zhang and Oliferenko, 2014) (yFS978) were shifted to the restrictive temperature of 35°C and sampled at 2, 4 and 6 hours. Cdc25-NeonGreen signal, GST-NLS-mCherry signal and cell length were measured microscopically in individual cells. The concentration of Cdc25 and GST-NLS-mCherry was calculated as the total nuclear fluorescent signal divided by the cell size. The mean signal from 25-cell bins are shown in large opaque symbols with error bars depicting standard deviation.

We next examined the dynamic range of Cdc25 size-dependent expression. We arrested cells in G2 with a temperature-sensitive allele of *cdc2* and measured the concentration of Cdc25 relative to Wee1 as cells grew from a normal size of around 15 μm to over three times that size. As cells grew, Cdc25 concentration increased linearly with size (Figure 1C), as has been seen for G2-and checkpoint-arrested cells (Moreno et al., 1990; Kovelman and Russell, 1996), showing that it is an accurate measure of cell size well beyond the normal length of G2.

To confirm the bulk analysis of Cdc25 concentration, we analyzed the expression of Cdc25-NeonGreen in individual cells. Because of the low level of Cdc25 expression, the Cdc25-NeonGreen data is noisy, precluding detection of the two-fold change in Cdc25 levels expected over a normal cell cycle. Nonetheless, as previously reported for Cdc25- GFP (Lu et al., 2012), Cdc25-NeonGreen concentration increases sufficiently when cells are arrested in G2 and allowed to grow to four times their normal size for an approximately four fold increase in Cdc25 concentration to be robustly measured (Figure 1D). As previously reported (Lopez-Girona et al., 1999; Lu et al., 2012), we observe that Cdc25 is predominantly nuclear in G2 (Figure S2). Both Cdc25 and Cdc2 shuttle in and out of the nucleus (Booher et al., 1989; Lopez-Girona et al., 1999). However, we assume that as Cdc25 concentration increases, it proportionally increases in all of its subcellular localizations.

We tested if the size-dependent expression of Cdc25 was regulated transcriptionally by measuring steady-state transcript levels in synchronized cell cultures. Mirroring protein levels, the concentration of *cdc25* transcript rises about 2 fold during G2 and then drops during mitosis, consistent with previous data (Moreno et al., 1990) (Figure 2A). Furthermore, we see a similar increase in *cdc25* transcript concentration at the single cell level (Figure 2B). *cdc25* transcript concentration, as assayed by single-molecule RNA-FISH (smFISH), increases linearly with cell size during G2 from a relative concentration of one at the beginning of the G2 to a relative concentration of two at the G2/M transition. It then drops back to one in post-mitotic cells, resetting the system for the next cell cycle.

**Figure 2:**
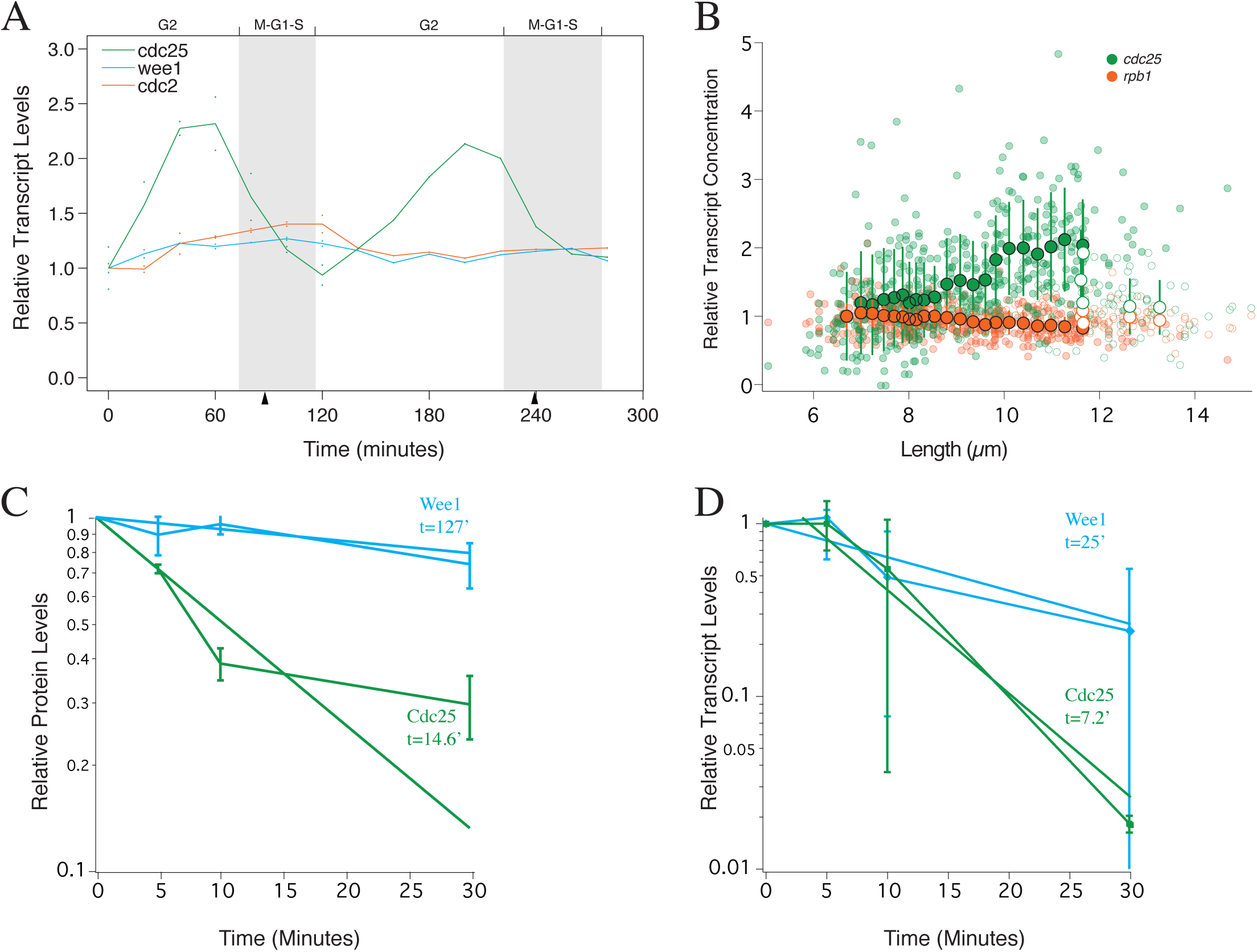
*cdc25* transcript is expressed in proportion to cell size. **(A)***cdc25* **transcript concentration doubles during G2.** Wild-type cell (yFS105) were elutriation synchronized in early G2 and followed through two synchronous cell cycles. Samples were taken every 20 minutes for RNA quantitation and examined microscopically for septation. Steady-state mRNA levels were determined using the NanoString nCounter method with custom probes and normalized to total mRNA counts. Data points represent independent biological replicates, the lines connect the mean of those points. The midpoint of septation for each cycle is marked with an arrowhead and the inferred M-G1-S phases of the cycles are indicated in gray. For the first two hours, n=2, for the rest of the time course, n=1. **(B)** *cdc25* transcript concentration increases with cell size in individual cells. Asynchronous wild-type cells (yFS105) were simultaneously analyzed for *cdc25* and *rbp1* transcript number by single-molecule RNA FISH. Data from individual cells is shown as small translucent symbols. Data from binucleate cells, which are in anaphase or G1, are shown as small open symbols. The mean transcript numbers from 50-cell bins of mononucleate (G2 and metaphase) cells are shown in large opaque symbols with error bars depicting standard deviation. Mean values from binucleate (anaphase, G1 and S-phase) cells are shown as large open symbols. **(C) Cdc25 protein is unstable.** Cells expressing Cdc25-Rluc and Ade4-Bluc (yFS874) were treated with 100 μg/ml cycloheximide and sampled as indicated for luciferase quantitation. As a comparison, cells expressing Wee1-Rluc and Ade4-Bluc (yFS810) were similarly assayed. The mean and standard error of the Ade4-nomalized Cdc25 and Wee1 signal, relative to time 0, is shown. n=3 for Cdc25; n=2 for Wee1. **(D)** *cdc25* **transcript mRNA is unstable.** Wild-type cells (yFS105) were treated with 15 µg/ml thiolutin and sampled as indicated for RNA quantitation by qRT-PCR. The mean and its standard error, relative to time 0, are shown. n=3.

We considered two explanations for the increase in the concentration of the *cdc25* transcript and its protein product during G2. The first explanation is that *cdc25* is turned on in early G2 and accumulates with pre-steady-state kinetics without reaching an expression equilibrium before cells enter mitosis. In this scenario, activation of the *cdc25* promoter in G2 creates a situation in which the rate of synthesis of *cdc25* mRNA is greater that the rate of its degradation, causing the *cdc25* mRNA to accumulate. In general, the rate of mRNA synthesis is of any gene is dependent on its rate of promoter initiation, but independent from the concentration of the mRNA produced, whereas the degradation rate of the same mRNA in dependent on its concentration (Wang et al., 2002). Thus, when a promoter is turned on, the mRNA expressed accumulates out of steady state (with a greater rate of synthesis than degradation) until the concentration of the mRNA increases to the point at which its degradation rate matches its synthesis rate, and steady-state expression is restored. The time it takes equilibrium to be reached is dependent on the half life of the protein and its mRNA. So, if Cdc25 and the *cdc25* mRNA were unusually stable, then Cdc25 could increase in concentration with pre-steady-state kinetics for all of G2. In such a model, the increase in Cdc25 concentration is time-dependent, not size-dependent.

The second explanation is that Cdc25 protein concentration is at a size-dependent steady state throughout G2, and thus serves as a direct measure of cell size. In this scenario, the rate of synthesis and degradation of the *cdc25* mRNA are balanced throughout G2, but that as the cell grows the rate of *cdc25* transcription increases or its degradation rate decreases, such that the level of *cdc25* transcript is size-dependent, leading to an increasing concentration of the *cdc25* mRNA and its protein product.

It is possible to distinguish between time-dependent pre-steady-state accumulation and size-dependent steady-state expression by examining the half-lives of the Cdc25 protein and its transcript. The time it takes a protein to come to equilibrium after an increase in transcription is determined by the half-life of the protein and its transcript (Mehra et al., 2003; Belle et al., 2006). Therefore, for Cdc25 to accumulate in pre-steady-state kinetics for the approximately 2 hour fission yeast G2 (or for the 6 hour arrest in Figure 1C), it would need to have transcript and protein half lives on the order of hours. On the contrary, we find that the half-life of Cdc25 protein is about 15 minutes (Figure 2C) and the half-life of the *cdc25* transcript is about 7 minutes (Figure 2D), consistent with previously reported data (Eser et al., 2016). These results demonstrate that Cdc25 levels do not increase in G2 due to pre-steady-state accumulation and supports a model in which Cdc25 expression increases at a size-dependent equilibrium.

Regulation of Cdc25 translation has also been proposed to be involved in size control (Daga and Jimenez, 1999). The *cdc25* transcript has an unusually long and structured 5' UTR, which makes it sensitive to the cells translational capacity, leading to the hypothesis that Cdc25 translation could be a readout of ribosome number and thus cell size (Daga and Jimenez, 1999). We tested the importance of this level of translational regulation for the size-dependent expression of Cdc25 by using a strain in which the *cdc25* 5' UTR is deleted (Daga and Jimenez, 1999). We find that the size dependent expression of Cdc25 does not require the 5' UTR of its transcript, showing that this mechanism of translational regulation is not necessary for size dependent expression (Figure S3A). Furthermore, the strain returns to its normal size rapidly after being elongated in a G2 arrest, demonstrating active cell size homeostasis (Figure S3B). Therefore, we proposes that, instead of playing a direct role in the mechanism of size homeostasis, translational control of Cdc25 may play a role in setting the size threshold in response to nutrient conditions (Fantes and Nurse, 1977) by modulating the amount of Cdc25 translated from size-dependent levels of the *cdc25* transcript.

Size control by size-dependent expression of an unstable mitotic activator has been proposed in a number of eukaryotic systems, including fission yeast, protists and mammalian cells (Miyamoto et al., 1973; Herring, 1974; Fantes et al., 1975; Polanshek, 1977; Tyson et al., 1979; Wheals and Silverman, 1982). A hallmark of this mechanism of size control is the phenomenon of excess mitotic delay, in which short pulses of the protein-synthesis inhibitor cycloheximide cause longer mitotic delays the closer they are applied to mitosis (Mitchison, 1971). These results have been interpreted in the context of the unstable-activator hypothesis (Wheals and Silverman, 1982; Tyson, 1983). This hypothesis posits that cell size is regulated by the size-dependent expression of an unstable mitotic activator, which triggers mitosis when it reaches a critical threshold in late G2. Since the activator rapidly decays during short G2 pulses of cycloheximide, a pulse in early G2 allows cells sufficient time to resynthesize the activator before mitosis, but a pulse applied later in G2 provides insufficient time for the activator to be resynthesized, thus delaying mitosis.

Fission yeast exhibit excess mitotic delay in response to cycloheximide pulses (Herring, 1974; Polanshek, 1977). Our results suggest that Cdc25 is an unstable activator that regulates cell size in fission yeast. To test if Cdc25 behaves as predicted by the unstable-activator model, we measured the kinetics of Cdc25 degradation and reaccumulation during and after a cycloheximide pulse. As predicted, Cdc25 levels fall during the pulse and then return to pre-pulse levels (Figure 3A). Importantly, the cycloheximide-treated cells do not divide until Cdc25 recovers to the level at which untreated cells divide (Figure 3A) and the delay in Cdc25 recovery matches the delay in mitotic entry (Figures 3A,B), suggesting that recovery of Cdc25 to a critical threshold is required to trigger the G2/M transition.

**Figure 3:**
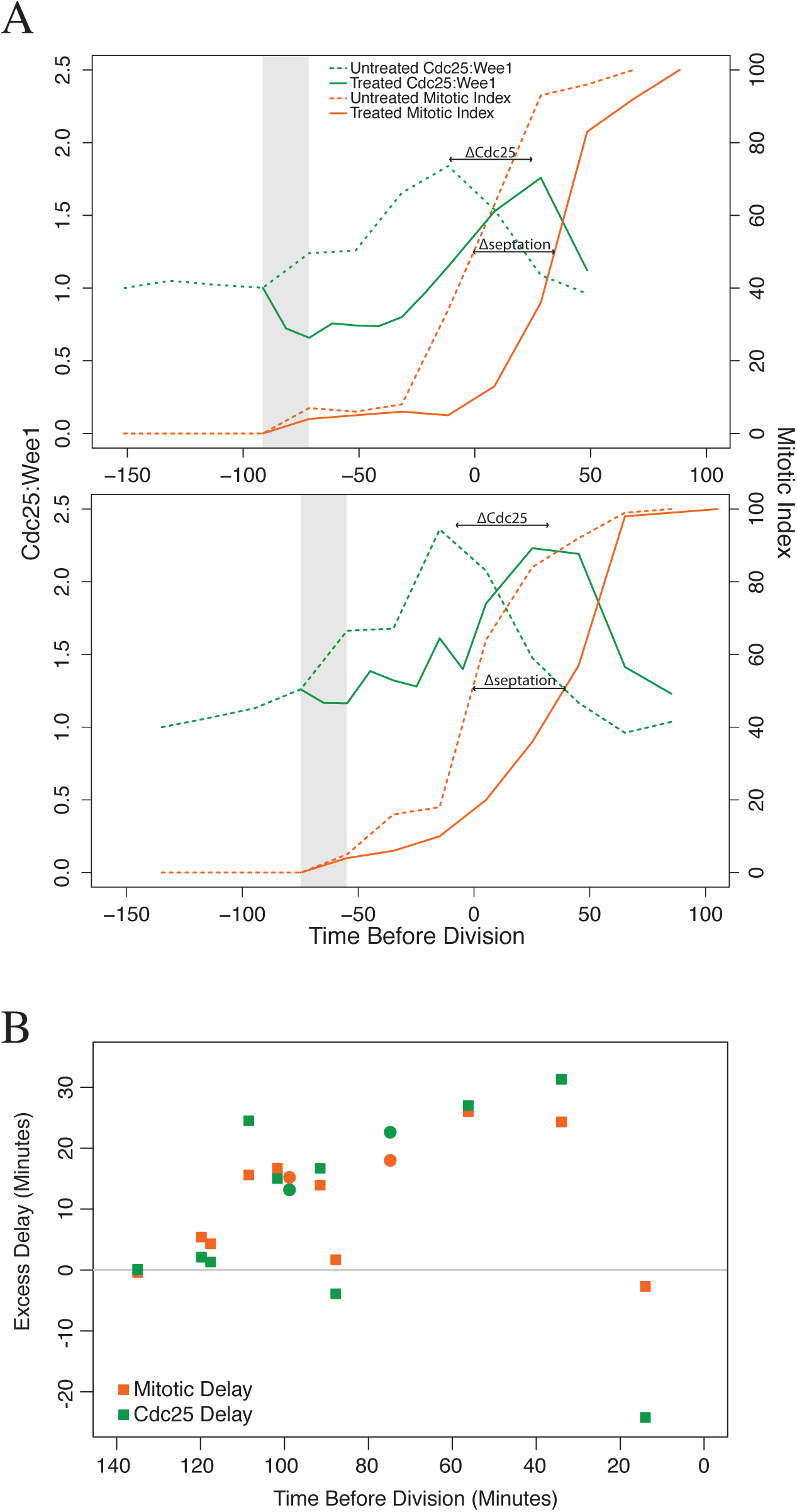
Cdc25 behaves as an unstable activator of mitosis. **(A) The delay in Cdc25 expression after a cycloheximide pulse mirrors the delay in mitotic entry.** Cells expressing Cdc25-Bluc and Wee1-Rluc (yFS870) were elutriation synchronized in early G2. Samples were taken every 20 minutes for luciferase quantitation and examined microscopically for septation. At the indicated times before division of the untreated cells, the culture was split and one half was treated with 100 μg/ml cycloheximide for 20 minutes. **(B) Quantitation of cycloheximide-induced delay in Cdc25 expression and mitotic entry.** Data from twelve experiments conducted as described in (A) is displayed. The experiments in (A) are shown as circles.

Our model makes specific predictions about the role of Cdc25 expression kinetics in triggering the G2/M transition. To test if these predictions are consistent with the detailed understanding of the G2/M regulatory network (Morgan, 2006), we integrated our hypotheses into a quantitative model of fission yeast cell-cycle dynamics (Novak and Tyson, 1995). We modified the model to include size-dependent increase in Cdc25 concentration and found realistic parameters under which such an increase was sufficient to drive stable cell cycles (Figure 4A) and to maintain size homeostasis (Figures 4B,C).

**Figure 4:**
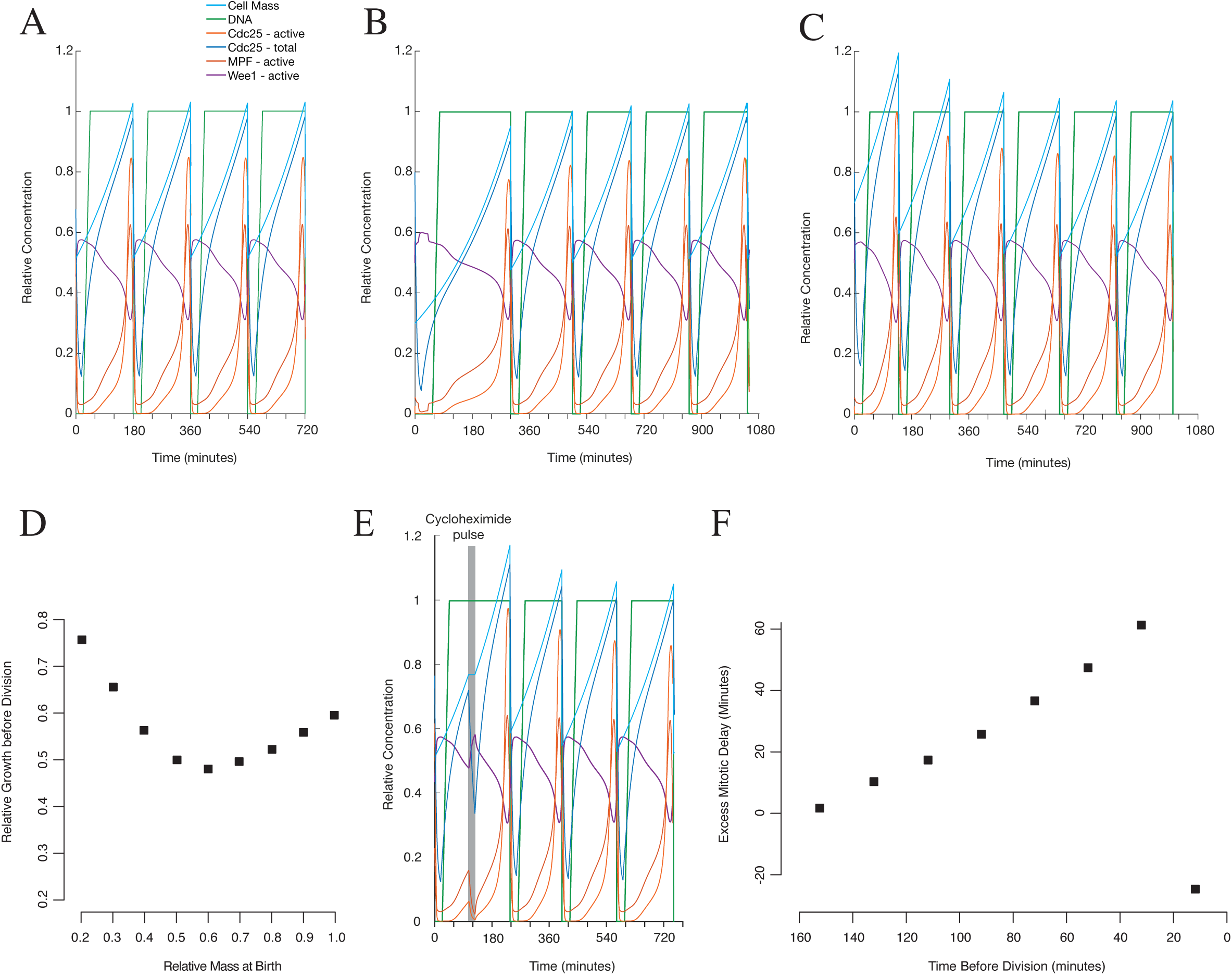
Mathematical modeling of cell-size control by Cdc25 expression. **(A) A Cdc25-concentration-regulated model of the cell cycle.** Simulation of the cell cycle using an ordinary differential equation model in which the size-dependent expression of Cdc25 triggers entry into mitosis at the appropriate size. **(B,C) The model maintains cell-size homeostasis. (B)** The cell cycle simulated as in (A), but initialized with a cell 60% the normal size at mitosis. **(C)** The cell cycle simulated as in (A), but initialized with a cell 140% the normal size at mitosis. **(D) Size at birth and growth during the cell cycle are negatively correlated.** The results of simulations such as those shown in (B) and (C) initiated at a range of cell masses are platted as the amount of mass (relative to normal size at division) added during the cell cycle against the mass of the cells at birth. Cells at size homeostasis are born at a mass of 0.5 and add a mass of 0.5 during their cell cycle. Active size control is depicted by the negative slope for cells up to a birth size of 0.6 (120% of normal). Cells born at larger sizes show a positive correlation because they grow for the minimum time possible and, during that minimum time, larger cells accumulate more mass. **(E) Simulation of cycloheximide-induced delay in Cdc25 expression and mitotic entry.** The cell cycle simulated as in (A), but with a simulated pulse of cycloheximide, during which the synthesis of all proteins and the increase in cell mass is set to 0, between 120 and 140 minutes (60 and 40 minutes before cell division would have happened without the pulse). Compared with (A), in which the cells divide at 180 minutes, division is delayed for about 40 minutes. **(F) Quantitation of cycloheximide-induced delay simulations.** The simulation in (D) was run with simulated cycloheximide pulses at various times from 20 to 140 minutes before cell division would have occurred in an untreated cell. For each simulation, the extent of cell cycle delay was recorded and plotted against time of the pulse.

To visualize the active size homeostasis in our model, we plotted size at birth versus growth during the cell cycle for cells simulated to be born at a variety of sizes (Figure 4D). A negative slope on such a graph is indicative of active size control, because small cell grow more before dividing than large cells (Fantes, 1977; Jun and Taheri-Araghi, 2015). Such a negative slope is observed up until a size at which the growth during a minimum-length cell cycle results in cells bigger than the normal size at division (Fantes, 1977). After that, the slope becomes positive because larger cells grow more during the minimum-length cell cycle than smaller cells. We see a negative slope diagnostic of active size control in simulated cells smaller than about 120% of normal size and then a transition to a positive slope in larger cells (Figure 4D), demonstrating functional size homeostasis in our mathematical model.

We then simulated the effect of cycloheximide pulses on the system and found that it recapitulated the excess delay phenomenon (Figure 4E), in agreement with our experimental data (compare Figures 3B and 4F). This model-based analysis demonstrates that size-dependent expression of Cdc25 provides a biochemically plausible mechanism for size control in fission yeast and accounts for the excess delay phenomena seen in fission yeast and other eukaryotes.

It should be noted that our mathematical model is designed to recapitulate the regulatory logic of the cell cycle not to quantitatively capture actual rate constants. Therefore, while our model qualitatively reflects results from this work and previous papers, it varies from such experimental results in kinetic details, such as the extent of excess delay in Figure 4F. However, since we have no data to constrain most of the biochemical rate constants in our model, optimizing them to fit experimental observations would not provide any biological insight.

Our data and analysis support a model in which size-dependent expression of the *cdc25* transcript leads to size-dependent expression of the Cdc25 mitotic inducer and thus size-dependent entry into mitosis. When cells are small, the activity of Cdc25 is insufficient to dephosphorylate and activate the Cdc2 CDK. When cells reach a critical size, the concentration of Cdc25 reaches the point at which it can begin to dephosphorylate Cdc2, which in turn hyper-activates Cdc25, leading to full dephosphorylation of Cdc2 and committing cells to mitosis (Rhind and Russell, 2012).

This model raises the question of how size-dependent regulation of Cdc25 levels interacts with other modes of Cdc25 regulation, such as translational regulation in response to nutritional conditions and post-translational regulation in response to stress and checkpoint signaling (Furnari et al., 1997; Shiozaki and Russell, 1997; Daga and Jimenez, 1999; Petersen and Nurse, 2007; Rhind and Russell, 2012). We propose that the size-dependent expression of the *cdc25* transcript is the mechanism that allows cells to divide at a particular size, but it is not the mechanism which regulates what that size is. It is the other modes of regulation that control the amount of Cdc25 activity produces from a given level of *cdc25* transcript, thereby determining the size a cell must attain before reaching the critical size threshold for division in response to environmental conditions.

This model also raises the question of how the *cdc25* transcript is maintained at a size-dependent level. The observation that doubling the *cdc25* gene dosage does not result in a dramatic reduction in cell size at division (Russell and Nurse, 1986 and our unpublished data) suggests that *cdc25* transcript levels are not solely determined by the rate of transcriptional initiation. This conclusion leads us to suspect that the stability of the *cdc25* transcript may be regulated in a size-dependent manner, although further work will be required to substantiate such speculation.

Because the Cdc25 phosphatase and its CDK substrates are well-conserved across fungi and metazoa (Morgan, 2006), size-dependent increase in concentration of Cdc25 provides a potentially wide-spread solution for the question of size control in eukaryotes. Although G1 size control mechanisms function in many eukaryotic species, including fission yeast (Amodeo and Skotheim, 2015), G2 size control may collaborate with G1 size control, as is evident in fission yeast and recently reported in budding yeast (Fantes and Nurse, 1978; Mayhew et al., 2017)

## Experimental Procedures

Bulk protein levels were determined by measuring the activity of translational fusions to beetle or *Renilla* luciferase expressed from the endogenous gene loci using a modification of the Dual-Luciferase Reporter Assay (Promega, Madison WI). Bulk steady-state mRNA levels were determined using the NanoString nCounter method and normalized to total mRNA counts. Transcript number in individual cells was determined by smFISH and normalized to transcript number in the smallest cells. mRNA half-lives were determined by qRT-PCR after inhibition of pol II in a synchronous G2 culture with 15 μg/ml thiolutin (Mendell et al., 2000). Mathematical cell-cycle models were implemented in MatLab (MathWorks, Natick MA) with custom code, available upon request.

## Acknowledgements

We are grateful to Jenny Benanti, Dan McCollum, Peter Pryciak, and members of the Benanti and Rhind labs for insightful suggestions on this work and constructive comments of the manuscript. We acknowledge the pioneering work in the Mitchison lab on the phenomenon of excess mitotic delay and thank Alan Herring for discussing his unpublished work on the subject. We thank Snezhana Oliferenko for the GST-NLS-mCherry construct and Bela Novak for help recreating his fission yeast cell cycle model. This work was supported by NIH GM098815 to NR and the Medical Research Council to SM. This work was performed under the auspices of the U.S. Department of Energy by Lawrence Livermore National Laboratory under contract DE-AC52-07NA27344 (LLNL-ABS-691918).

## Author Contributions

Conceptualization: DK,NR; Methodology: DK,XS,ED,MO,MBM,SM,NR; Software:

ED,MBM; Formal Analysis: DK,MBM,SM,NR; Investigation: DK,XS,ED,MO,MM,AH,MBM,NR; Data Curation: NR; Writing–Original Draft: NR; Writing-Review & Editing: DK,XS,ED,MO,MM,AH,MBM,SM,NR; Visualization: DK,MBM,SM,NR; Supervision: SM,NR; Project Administration: NR; Funding Acquisition: SM,NR

## Supplemental Experimental Procedures Strain Construction and Maintenance

Strains were created and cultured using standard techniques (Forsburg and Rhind, 2006).

Cells were grown in yeast extract plus supplements (YES) at 30°C, unless otherwise noted. Strains with temperature-sensitive alleles were grown at 25°C for permissive growth and switched to 35°C for non-permissive growth. The follow in strains were used.

yFS105 h- leu1-32 ura4-D18 (14.3 +/- 1.0 μm)
yFS810 h-leu1-32 ura4-D18 ade4-Bluc (KanMX) wee1-Rluc (NatMX) (15.2 +/- 1.3 μm)
yFS870 h-leu1-32 ura4-D18 wee1-Rluc (NatMX) cdc25-Bluc (KanMX) (15.7 +/- 1.2 μm)
yFS874 h-leu1-32 ura4-D18 ade4-Rluc (NatMX) cdc25-Bluc (KanMX) (12.9 +/- 0.7 μm)
yFS885 h+leu1-32 ura4-D18 ade4-Rluc(NatMX) cdc25-d1-Bluc (KanMX) (13.8 +/- 1.1 μm)
yFS893 h-leu1-32 ura4-D18 cdc2-L7 wee1-Rluc (NatMX) cdc25-Bluc (KanMX) yFS971 h-leu1-32::nmt81-GST-NLS-mCherry ura4-D18 cdc2-33 cdc25-NeonGreen (HphMX)
yFS982 h-leu1-32 ura4-D18 cdc2-L7 ade4-GFP (HphMX)

The average size at septation (+/- standard deviation) for selected strains show that the luciferase-tagged Cdc25 and Wee1 proteins are functional, causing less that a 10% change in average length at septation.

The following primers and plasmid templates were used to create integration cassettes.

| Cassette | Primers | Plasmid | Plasmid reference |
| --- | --- | --- | --- |
| ade4-Bluc (KanMX) | DK26 DK27 | pFS345 | this work |
| wee1-Rluc (NatMX) | NR186 NR187 | pFS346 | this work |
| cdc25-Bluc (KanMX) | DK6 DK7 | pFS345 | this work |
| cdc25-NeonGreen (HphMX) | MM6 MM41 | pFA6-mNG:hyg | (Willet et al., 2015) |
| ade4-GFP (HphMX) | MO294 MO296 | pFA6a-GFP-hph | (Sato et al., 2005) |

DK6 TTGGCCAAAGTGTGTTAGCTTCCCCAGACGTTAATGATTCTCCTACTGCC ATGCATTCCCTCTCTACACTTAGAAGATTTCGGATCCCCGGGTTAATTAA DK7 AGAAAAAACTTAGGTTTAGAAAGTTGAATATATAAGAGTATACTTCAGGCTAGGTAAAGTATTGAGTCAGCCTAAAATCAGAATTCGAGCTCGTTTAAACDK26 TTGCAGAAGATGAGGAACGTGAAGCTCCCGAAGACATTTCTCTCCATAACACACATTCAGATGTTACTTTTGATTTTGTTCGGATCCCCGGGTTAATTAA DK27 GATTAGAGCATCAATCTAGACAAAGTAAATGGAGGATTGGTTATTATAATAAAGCACTAAGCATTGAATAAATT GGGGAAGAATTCGAGCTCGTTTAAACNR186 CCATAATTTATGAAGGTATTCATGGATCTTCTTCTAACCCCCAGGGTG ATCAAATGATGGAAGATTGGCAGGTGAATGTTCGGATCCCCGGGTTAATTAA NR187 GCTAAACAGATTTTGGAAGCCATTCCCTTATTTCGCAATTTCGCAGTAATAAACATTGAGAACAAGAGTCTCTAAAAGGTGAATTCGAGCTCGTTTAAAC MM6 AGAAAAAACTTAGGTTTAGAAAGTTGAATATATAAGAGTATACTTCAG GCTAGGTAAAGTATTGAGTCAGCCTAAAATCACGCACTTAACTTCGCATCTG MM41 TTGGCCAAAGTGTGTTAGCTTCCCCAGACGTTAATGATTCTCCTACTGCCATGCATTCCCTCTCTACACTTAGAAGATTTCGGATCCCCGGGTTAATTAACAT MO294 GATTAGAGCATCAATCTAGACAAAGTAAATGGAGGATTGGTTATTATAA TAAAGCACTAAGCATTGAATAAATTGGGGAATTATTCCTTTGCCCTCGGACGAGT MO296 TTGCAGAAGATGAGGAACGTGAAGCTCCCGAAGACATTTCTCTCCATAACACACATTCAGATGTTACTTTTGATTTTGTTAGTAAAGGAGAAGAACTTTTCACTG

Plasmids and plasmid sequences created for this work are available at AddGene <http://www.addgene.org>.

### Synchronization and Time Course

Cells were synchronized by centrifugal elutriation in a Beckman JE-5.0 elutriating centrifuge rotor (Willis and Rhind, 2011). Time points were taken every 20 minutes to measure septation and for protein samples. Septation was monitored by counting unseptated, septated, and undivided pairs. Mitotic index was calculated as the ratio of septated and undivided pairs divided by total count for that time point. For the luciferase assay, samples were washed with cold water, pelleted and frozen in liquid nitrogen.

### Luciferase Assay

Cell pellets were processed for the luciferase assay following a modified procedure based on the Dual-Luciferase Reporter Assay (Promega, Madison WI). 5-10 OD pellets were lysed at 4°C in 200 μl 1X Passive Lysis Buffer by bead beating to a point where ~80% of the cells were lysed. Lysates were cleared at 16,000g at 4°C. 10 µl of cleared lysate was loaded per well in an opaque 96-well plate and each sample was read in triplicate at room temperature. For each well, 50µL of Luciferase Assay Substrate and Stop and Glow Buffer are added sequentially to assay for beetle followed by *Renilla* luciferase. After the addition of each substrate, the samples were equilibrated for 2 seconds followed by a 10 second measurement for luminescence.

### Cdc25 Quantitation by Fluorescent Microscopy

yFS978 cells were grown in EMM2-LUAH media at 25°C to mid log phase.

Asynchronous cells were shifted to 35°C, sampled after 2, 4 and 6 hours, fixed with 100% methanol at −20°C, mixed in equal proportions, rehydrated in 1x PBS and imaged on an DeltaVision OMX microscope with a 60x/1.42 NA objective and InSightSSI solid-state fluorescence illumination, which provides uniform illumination, facilitating accurate quantitation of the fluorescent signal. Wide-field images were acquired and manually analyzed using ImageJ 1.49q (Schneider et al., 2012).

### Transcript Quantitation

For NanoString quantitation, 1x10^7^ cells were fixed with 70% methanol and stored at - 80°C in 1 ml of RNALater (Ambion). For processing, cells were pelleted, resuspended in 600 μl RLT buffer (Qiagen) with 1% β-mercaptoethanol and lysed by bead beating. 200 µl of lysate was cleared at 16,000 g and 3 μl of supernatant was processed on a NanoString nCounter (Seattle, WA) with a custom code set according to the manufacturers instructions.

### Single Molecule RNA Fluorescence In Situ Hybridization (smFISH)

smFISH samples were prepared according to a modification of published protocols (Trcek et al., 2012; Heinrich et al., 2013). Briefly, cells were fixed in 4% formaldehyde and the cell wall was partially digested using Zymolyase. Cells were permeabilized in 70% EtOH, pre-blocked in BSA and salmon sperm DNA, and incubated over-night with custom Stellaris oligonucleotides sets (Biosearch Technologies) designed against cdc25 (CAL Fluor^®^ Red 610) and rpb1 (Quasar^®^ 670) mRNAs. Cells were mounted in ProLong Gold antifade reagent with DAPI (Life Technologies) and imaged on a Leica TCS Sp8 confocal microscope, using a 63x/1.40 oil objective. Optical z sections were acquired (z-step size 0.3 microns) for each scan to cover the depth of the cells. Cell boundaries were outlined manually and single mRNA molecules were identified and counted using the FISH-quant MATLAB package (Mueller et al., 2013). Cell area, length and width were quantified using custom-made ImageJ macros. The FISH-quant detection technical error was estimated at 6-7% by quantifying rpb1 mRNAs simultaneously with two sets of probes labeled with different dyes.

### Transcript Half Life and RT-qPCR

For calculation of transcript half-life, log phase cultures were treated with 15 μg/ml thiolutin to inhibit polymerase II and 10 OD samples were taken at 0, 5, 10 and 30 minutes. Samples were pelleted and frozen in liquid nitrogen. Total RNA was isolated from pellets using the Direct-zol kit (Zymo Research, Irvine, CA). First strand synthesis was performed using random hexamers and SuperScript III first strand synthesis kit (Invitrogen). qPCR was performed using Kappa SYBR Fast qPCR kit (Wilmington, MA). Transcripts were normalized to 0 time point and *srp7* as an internal control for a stable transcript. Primers for each target are as follows:

*cdc25* - ATGACCTGCACCAAGGCTAT, TCATTAACGTCTGGGGAAGC
*wee1* - GATGAGGTTTGCTGGGTTGA, CATTCACCTGCCAATCTTCC
*cdc13* - ACCACGAGCTGTCCTTAACC, TGCTTAACCGACCAGGTTCC
*upf2* - ATCCGCCAAAGCGTGGTATC, AAGCGCACTAAGCAGACGAG
*srp7* - GTGCATGTTCGGTGGTCTCG, AAGACCCGGTAGTGATGTGC.

Half life data was fit with exponential decay curves using Igor Pro (WaveMetrics).

### Protein Half Life

To measure protein half lives, strains with a luciferase-tagged protein of interest were grown to log phase, 100 μg/ml of cycloheximide was added and 10 OD samples were taken at 0, 5, 10 and 30 minutes. Samples were pelleted, frozen in liquid nitrogen and processed as described above for luciferase measurement. Half life data was fit with exponential decay curves using Igor Pro (WaveMetrics).

### Excess Delay Assay

To assay for excess delay, an elutriation time course, described above, was modified by splitting the synchronized culture into two subcultures. One subculture was treated with a 20 minute pulse of 100 μg/ml of cycloheximide. Cycloheximide was removed by filtration and cells were put into fresh media and sampled every 20 minutes for septation and luciferase activity.

### Replicating and modifying the Novak and Tyson fission yeast cell cycle model

A previously published model of the fission yeast cell cycle, on which we based our work and which we refer to as NT95, consists of 18 differential equations and ~50 rate constant parameters (Novak and Tyson, 1995). We refer interested readers to that article for mathematical details of the full ODE model. In the model, cell size drives the inhibitory phosphorylation of Wee1 (via PK and Nim1). Thus, as the cell grows over the course of the cell cycle and active MPF levels rise, Wee1 is kept in its inactive state. In NT95, Cdc25 concentration is not size-dependent. It is also worth noting that G1/S progression is modeled by a “black box” automata in which certain rate constant parameters are set to different values depending on whether the cell has reached a certain size or on whether a certain amount of time has elapsed since division. Again, we refer readers to NT95 for details of this aspect of the model. As the G2/M transition is the more important point for fission yeast size control, we retained this automata model for G1/S progression. Our replication of NT95 was fully implemented in MATLAB (code available upon request). We obtained initial conditions by simulating from the model with the growth rate set to 0, taking the values of all species once they appeared to equilibrate (after ~5 cycles). Rate constants were taken directly from NT95. Cell growth was assumed to be exponential with a mass doubling time fixed at 180 minutes. Simulations were generated with MATLAB’s ode15s solver (variable order, multistep) for stiff systems of ODEs.

To test that the model had been successfully replicated, we recreated Table 1 from NT95, simulating from our version of NT95 and estimating the proportion of time spent in each part of the cell cycle under the 21 different genetic conditions tested. All our estimates corresponded exactly to those in Table 1 of NT95.

### Modifying and fitting the wild-type fission yeast cell cycle model

We modified NT95 in two ways: 1) by removing the size dependence from the Wee1 edges of the biochemical network, instead making total Cdc25 concentration dependent on cell mass; 2) by removing the rate equation and parameters for species X (an arbitrary species sensitive to unreplicated DNA introduced into the model to mediate mitotic progression). On this second modification, we replaced the rate equation of species W, the downstream target of X, with the dynamics of species X. We made this modification after noticing that the functional role of species W paralleled that of Cds1 in the fission yeast mitotic network, and consequently, required faster and more immediate decay dynamics following DNA replication. Species W is now named ‘Cds1’ in our model, and with the change in dynamics, total Cds1 is not capped at 1 as species W is in NT95. The two rate equations we introduced into our modified model were:

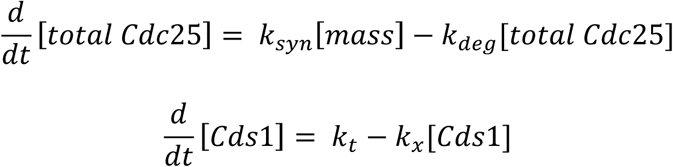

To estimate rate constants for the wild-type model, we first simulated for three cycles from the original NT95 model. We then treated these simulated curves for each species as data. Assuming that the rate constants for the modified model, which we refer to as SC16, would not be too far removed from their previous values, we used a direct, pattern search optimization routine to estimate rate parameters for our wild-type SC16 model. We used the sum of squared errors between three-cycle simulations from our SC16 model and the three-cycle simulations generated by the NT95 model as our objective function. All species were compared in this optimization. To constrain the optimization, we used lower bounds of 0 and upper bounds of 10 times the NT95 values of each rate parameter. The values of nearly every rate constant in the SC16 model remained unchanged from their NT95 values. Estimates for rate constants in the SC16 model that differed are shown below.

| Parameter | Description | NT95 Value | SC16 Value |
| --- | --- | --- | --- |
| $k_3$ | Formation of unphosphorylated MPF | 10.0 | 22.875 |
| $k_{CAK}$ | Phosphorylation of Cdc2 threonine | 1.0 | 1.5 |
| $k_a$ | Phospho-activation of Cdc25 | 0.5 | 0.5625 |
| $k_{f1}$ | Dephosphorylation (N-terminal) of Wee1 | 1.0 | 3.1875 |
| $k_{e1}$ | Phosphorylation (N-terminal) of Wee1 | 1.0 | 4.2539 |
| $k_{f2}$ | Dephosphorylation (C-terminal) of Wee1 | 1.0 | 1.1875 |
| $k_{e2}$ | Phosphorylation (C-terminal) of Wee1 | 1.0 | 0.875 |
| $k_x$ | Degradation rate of Cds1 (species X in NT95) | 0.1 | 0.1313 |
| $k_i$ | Phosphorylation of intermediary enzyme (species IE) | 0.2 | 0.2625 |
| $V_{mik}''$ | Component of $k_{mik}$ parameter for MPF phosphorylation | 0.2 | 0.202 |
| $F$ | Multiplier of phosphorylated TA species for DNA replication | 15.0 | 142.877 |
| $F_b$ | Multiplier of W for Cdc25 deactivation | 2.0 | 1.998 |
| $k_{deg,dn}$ | Degradation of Cdc25 after initial 20 minute period of high post-mitotic degradation | 0.04 | 0.038 |

### Simulating cycloheximide-pulse experiments

To simulate the cycloheximide-pulse experiments, we introduced a new terminal event representing the pulse to interrupt the ODE solver. The two parameters of the pulse were the time post-G2 entry (in minutes) at which the pulse occurred and the duration of the pulse. The duration of the pulse was fixed to 20 minutes in all simulations while we varied the start time of the pulse from 0 to 120 minutes post-G2 entry (by 20 minutes). At the onset and for the duration of the pulse, the Cdc25 synthesis rate constant (k_syn_), the Cds1 synthesis rate constant (k_t_), the mass growth rate (μ), the Cdc13 synthesis rate constant (k_1_,_AA_), and the Mik1 synthesis rate constant (k_p_) were set to 0.0. All rate constants were restored to their original values after the pulse. We generated one pulse per simulation, and we recorded effects of the pulse on cell-cycle duration and mass at division.

---

**Figure S1:**
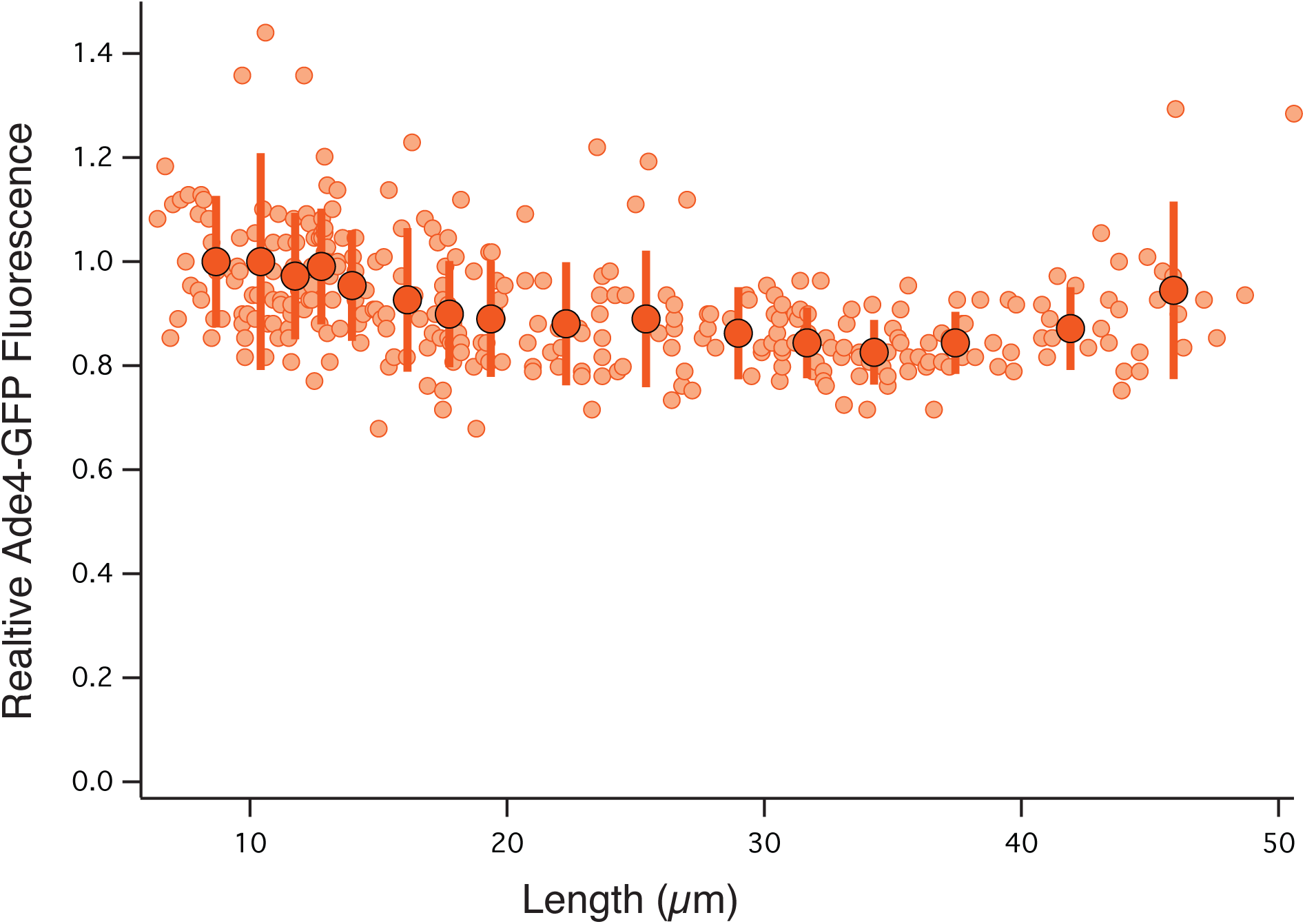
Ade4 maintains a constant concentration, independent of size. *cdc2-ts* cells expressing Ade4-GFP (yFS982) were shifted to the restrictive temperature of 35°C and sampled at 0, 2, 4 and 6 hours. Ade4-GFP signal and cell length were measured microscopically in individual cells. The concentration of Ade4 was calculated as the mean cellular fluorescence. The mean signals from 25-cell bins are shown in large opaque symbols with error bars depicting standard deviation.

**Figure S2:**
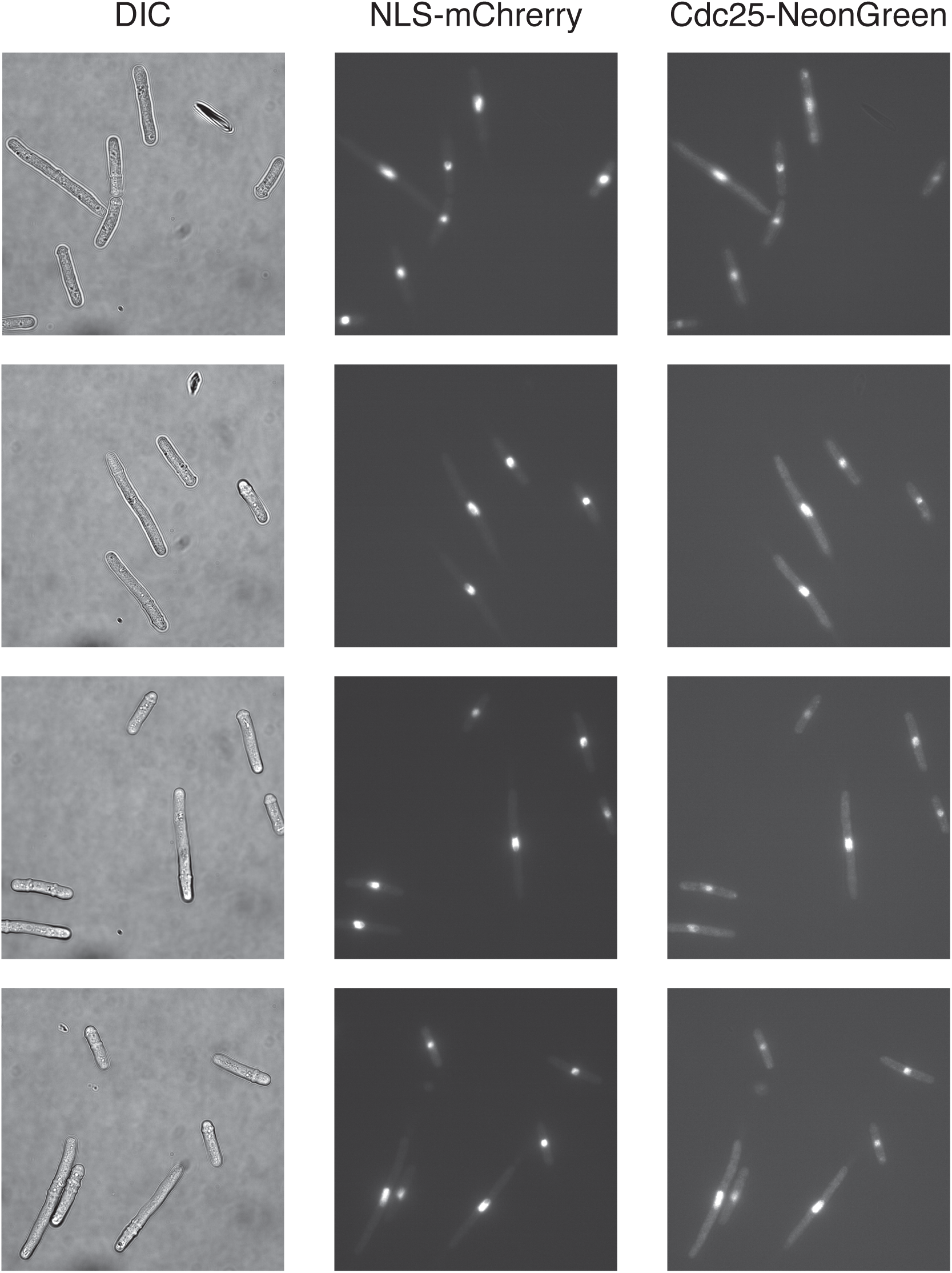
Images of fluorescent cells quantitated in Figure 1D. *cdc2-ts* cells expressing Cdc25-NeonGreen and GST-NLS-mCherry (yFS978) were shifted to the restrictive temperature of 35°C for 2, 4 or 6 hours, mixed and imaged by DIC and wide-field epifluorescence microscopy.

**Figure S3:**
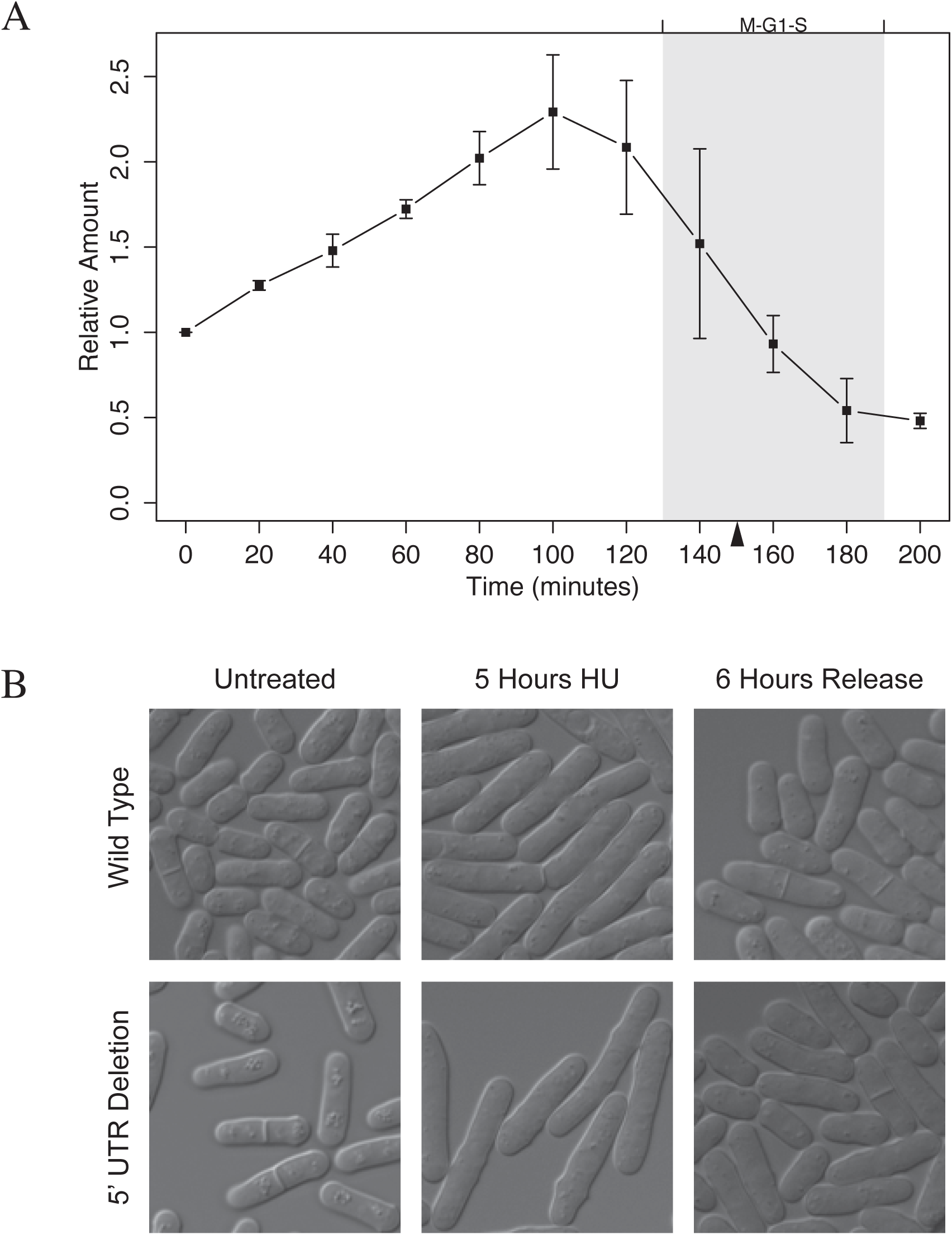
Size-dependent expression of Cdc25 does not require its 5’ UTR. **(A)**. Cells deleted for the *cdc25* 5’-UTR (Daga and Jimenez, 1999) and taged with luciferase (yFS885) were analyzed as in Figure 1A. **(B)**. Wild-type (yFS105) and cells deleted for the *cdc25* 5’-UTR (Daga and Jimenez, 1999) and taged with luciferase (yFS885) were arrested for 5 hours in 10 mM HU and released for 6 hours. Both strains rapidly return to their normal cell size after G2 elongation, demonstrating size homeostasis.

